# On-chip human lymph node stromal network for evaluating dendritic cell and T-cell trafficking

**DOI:** 10.1101/2023.03.21.533042

**Authors:** Brian J. Kwee, Adovi Akue, Kyung E. Sung

## Abstract

The lymph node paracortex, also known as the T-cell zone, consists of a network of fibroblastic reticular cells (FRCs) that secrete chemokines to induce T-cell and dendritic cell trafficking into the paracortex. To model the lymph node paracortex, we utilized multi-channel microfluidic devices to engineer a 3D lymph node stromal network from human cultured FRCs embedded in a collagen I-fibrin hydrogel. In the hydrogel, the FRCs self-assembled into an interconnected network, secreted the extracellular matrix proteins entactin, collagen IV, and fibronectin, as well as expressed an array of immune cell trafficking chemokines. Although the engineered FRC network did not secrete characteristic CCR7-ligand chemokines (i.e. CCL19 and CCL21), human primary TNF-α matured monocyte-derived dendritic cells, CD45RA^+^ T-cells, and CD45RA^-^ T-cells migrated toward the lymph node stromal network to a greater extent than toward a blank hydrogel. Furthermore, the FRCs co-recruited dendritic cells and antigen-specific T-cells into the lymph node stromal network. This engineered lymph node stromal network may help evaluate how human dendritic cells and T-cells migrate into the lymph node paracortex via CCR7-independent mechanisms.

## Introduction

The lymph node paracortex, otherwise known as the T-cell zone, is the region of the lymph node where dendritic cells (DCs) activate and present antigen to T-cells (*1*). The paracortex consists of podoplanin positive (PDPN^+^) fibroblastic reticular cells (FRCs), which ensheath an extracellular matrix rich conduit network that delivers antigens and other cytokines to lymphocytes in the T-cell zone (*1-3*). FRCs *in vivo* have been shown to secrete a variety of cytokines, including CCL19, CCL21, and CXCL4, which regulate the trafficking of DCs and T-cells, as well as IL-7, which influences the homeostasis of naïve T-cells (*1, 4, 5*). Furthermore, FRCs and the conduit network also serve as a cellular and molecular network that immune cells, including DCs and T-cells, migrate along and adhere to (*3, 6*). To date, our mechanistic understanding of the lymph node paracortex and fibroblastic reticular cells has mostly relied on *in vivo* murine models (*1-6*). However, it is not well known how accurately these models recapitulate human lymph node function, considering the fundamental differences between human and mouse immunology and lymphatics (*7, 8*).

Several microphysiological systems of the lymph node and T-cell zone have been utilized to model and study lymph node function *in vitro* or *ex vivo*. Murine FRC lines cultured on tissue culture polystyrene (TCPS) and on 3D nylon scaffolds coated with laminin have demonstrated the importance of lymphocyte-derived cytokines, in particular TNF-α and lymphotoxin-α/β, on ERTR-7 reticulum secretion by FRCs (*9*). Furthermore, primary murine FRCs cultured on macroporous polyurethane scaffolds with collagen I and Matrigel revealed how interstitial fluid flow upregulates CCL21 expression in FRCs (*10*). Other models of the lymph node, including microfluidic lymph node follicles (*11*), tonsil organoids (*12*), tonsil slices (*13*), and lymph node slices (*14*) have been developed to model humoral immunity and vaccine responses *ex vivo* and *in vitro*. However, 3D human models of the lymph node paracortex have been lacking, as the majority of studies utilizing human FRCs have evaluated their function on 2D TCPS (*13, 15, 16*).

In the present work, we engineered 3D, multi-channel microfluidic systems that model the human lymph node paracortex. We encapsulated human cultured FRCs in a collagen I-fibrin composite hydrogel and evaluated the ability of the FRCs to self-assemble into a 3D stromal network that mimics the lymph node paracortex. This network was found to secrete extracellular matrix proteins and an array of chemokines implicated in lymphocyte migration. Furthermore, we studied the ability of the FRC network to regulate the migration of co-cultured human dendritic cells and/or T-cells toward the network. This work demonstrates a 3D model that could potentially evaluate the ability of different human dendritic cells and T-cells to migrate toward the lymph node paracortex.

## Results

### Engineering the lymph node stromal network

To fabricate the 3D lymph node stromal network, commercially available human lymphatic fibroblasts from ScienCell™ were expanded and fluorescence activated cell sorting was used to enrich for CD45^-^CD31^-^PDPN^+^ FRCs, which were 99% positive for podoplanin after further expansion (Fig. S1). The expanded FRCs were encapsulated in a collagen I-fibrin composite hydrogel and then injected into Channel 5 of a custom designed, 6-channel polydimethylsiloxane microfluidic device at day -4 (Fig. 1). Blank collagen I-fibrin hydrogels were injected into Channels 2 and 4, while complete FRC media was injected into the remaining channels (Fig. 1B,C). After 4 days of culture, the FRCs spread and formed an interconnected, 3D cell network throughout the hydrogel in Channel 5 (Fig. 1C&2A, Fig. S2A,B). The FRCs also migrated and spread into the adjacent hydrogel in Channel 4 over 4 days (Fig. 2A, Fig. S3A,B). The extracellular matrix proteins entactin, collagen IV, and fibronectin were detected by immunohistochemistry in the surrounding hydrogel, adjacent to the FRCs after 4 days of culture in both Channel 4 and 5 (Fig. 2B,C). Collagen III, collagen VI, vitronectin, tenascin C, and laminin were not detected in the surrounding hydrogel.

**Fig. 1.**
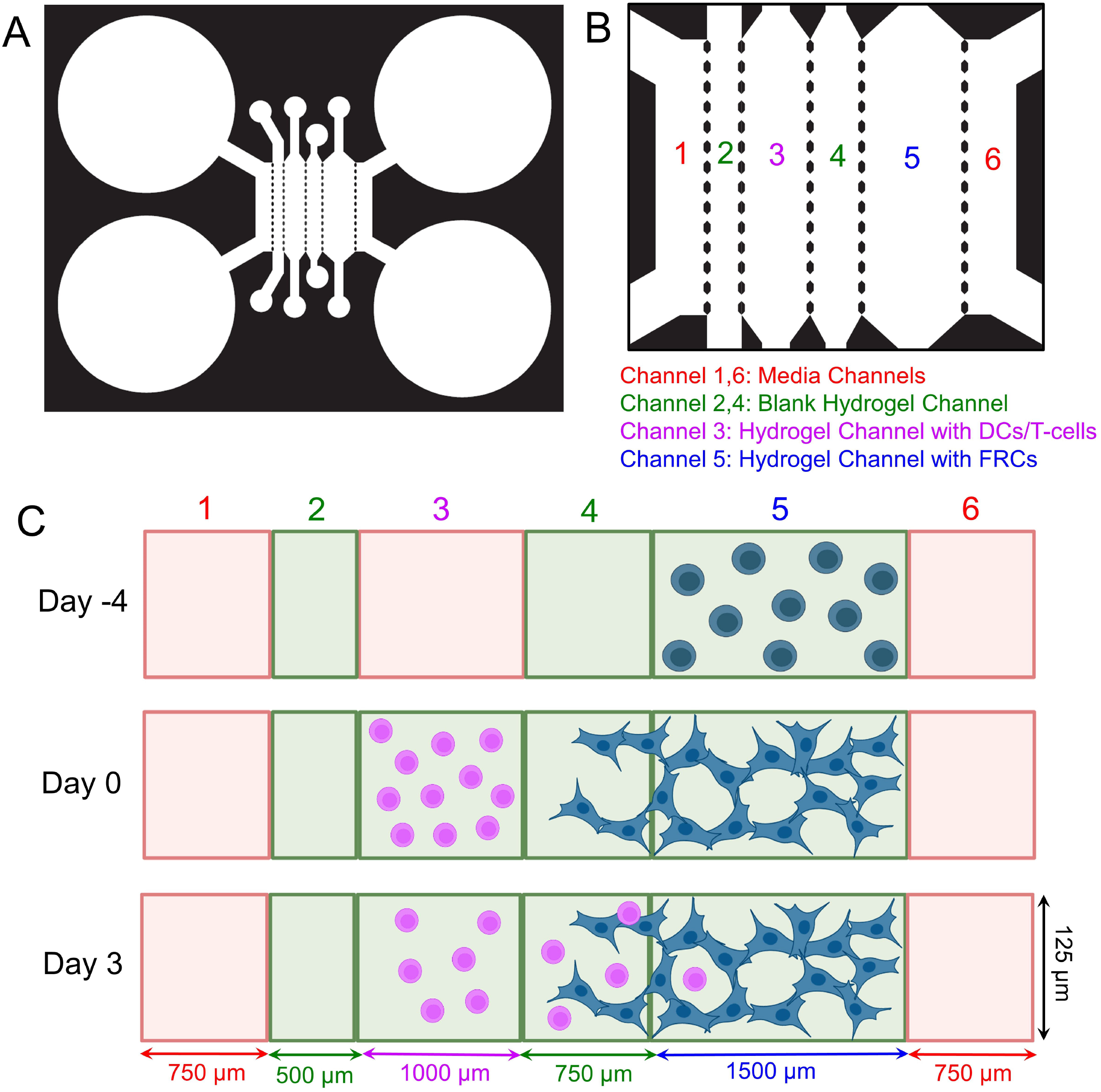
Engineering an on-chip lymph node stromal network. (A) Photolithography mask of microfluidic device with 6 fluid channels for fabricating the lymph node stromal network chip. Zoomed in image of photolithography mask of 6 channel device, denoting channel numbers and channel contents. (C) Graphical timeline of lymph node stromal network chip set up. At day -4, a blank hydrogel is injected into Channels 2 & 4, and a fibroblastic reticular cell (FRC) hydrogel (or a blank hydrogel) was injected into Channel 5. Media was added to Channels 1, 3, and 6. After 4 days of culture, a hydrogel loaded with dendritic cells or T-cells was injected into Channel 3. Migration of immune cells across channel 4 and into channel 5 was evaluated over the course of 3 days.

**Fig. 2.**
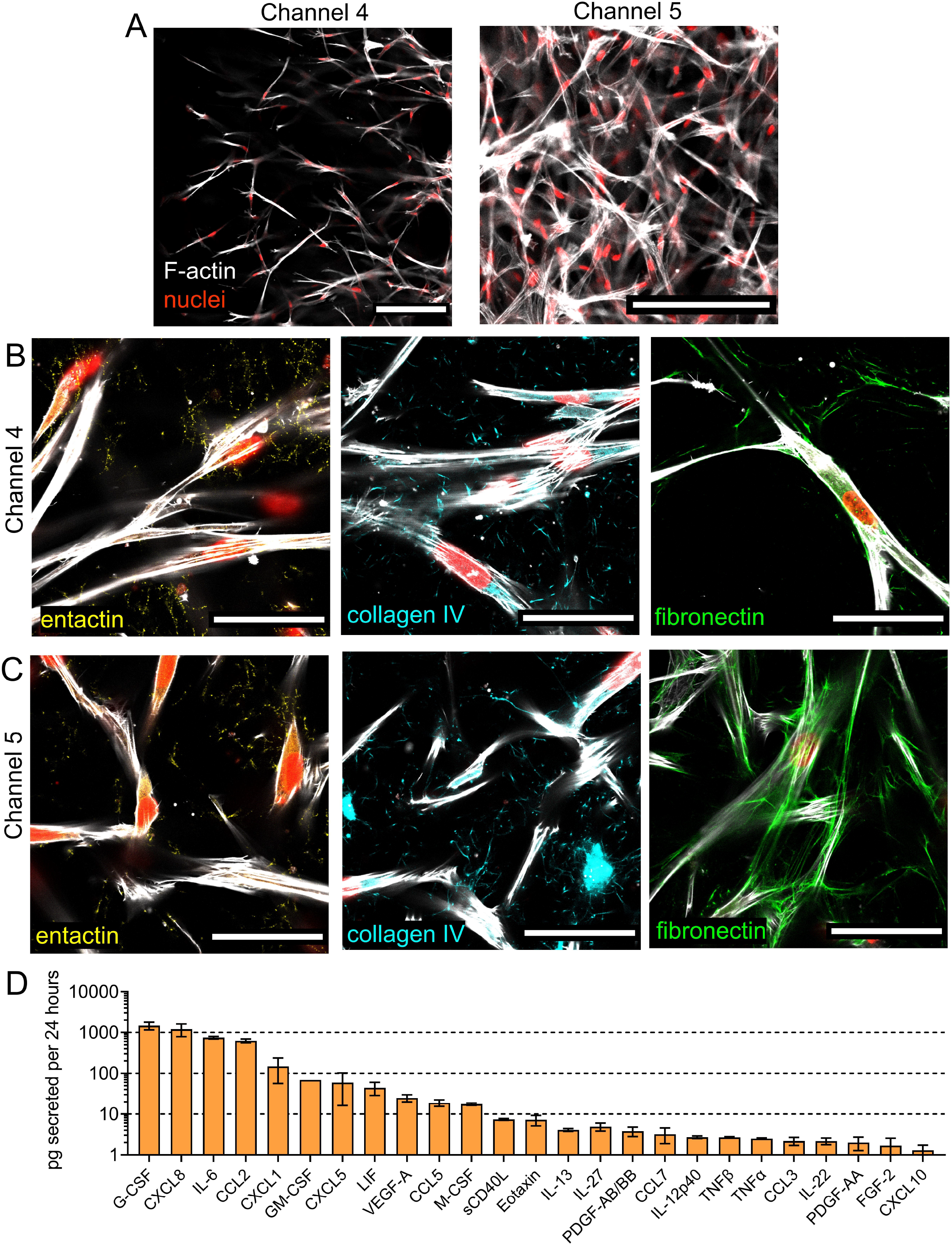
Lymph node stromal network characterization. (A) Representative confocal microscopy images of fibroblastic reticular cells (FRCs) in the collagen I-fibrin hydrogel at day 0 in Channel 4 and Channel 5 of the microfluidic device. Scale bar = 200 μm. (B) Representative confocal microscopy images of entactin, collagen IV, and fibronectin expression by FRCs on day 0 in Channel 4 and Channel 5. Red = nuclei, white = phalloidin F-actin, yellow = entactin, cyan = collagen IV, green = fibronectin, scale bar = 200 μm. (D) Picograms of chemokines and cytokines secreted by fibroblastic reticular cells over 24 hours from day 0 to 1. n = 4 devices. Data are presented as means ± SD.

Upon forming an interconnected FRC network at day 0, the culture media was switched to immune cell media. After 3 days of culture, the FRC network in Channel 4 and 5 showed minimal changes in structure and distance migrated (Fig. S2A, Fig. S3A). Quantitative analysis of 71 different cytokines and chemokines were performed on the conditioned media in the devices over the course of 3 days. Analysis of the 25 most upregulated chemokines and cytokines revealed the presence of chemokines previously implicated in dendritic cell migration and trafficking, including CCL2 (*17*), GM-CSF (*18*), CCL5, CCL7, and CCL3 (*19*) (Fig 2D, Fig. S4). Furthermore, the presence of chemokines and cytokines associated with T-cell migration and trafficking were detected, including CXCL8 (*20*), IL-6 (*21*), CCL2 (*22*), CCL5 (*23*), CCL7 (*24*), CCL3 (*25*), and CXCL10 (*26*) (Fig. 2D, Fig. S4). The cytokines CCL21, CXCL4, and IL-7 were not detected in the conditioned media.

### Migration of TNF-α matured monocyte-derived dendritic cells toward lymph node stromal network

The ability of TNF-α matured monocyte-derived dendritic cells (TNF-moDCs) to migrate toward the lymph node stromal network was then evaluated. CM-Dil labeled TNF-moDCs were encapsulated in a collagen I-fibrin hydrogel and injected into Channel 3 (Fig. 1B,C). TNF-moDC chemotaxis over 3 days was compared between devices with a blank hydrogel injected into Channel 5 (blank hydrogel) versus devices with a FRC-loaded hydrogel in Channel 5 (FRC hydrogel). TNF-moDC migration was quantified across Channel 4 and into Channel 5 (Fig. S5). At day 1, there were no significant differences in TNF-moDC migration across Channel 4 and into Channel 5 in the blank hydrogel compared to the FRC hydrogel (Fig. 3C). At day 2, TNF-moDCs migrated a significantly greater distance into Channel 5 of the FRC hydrogel compared to the blank hydrogel (Fig. 3D). At day 3, the number of migrating TNF-moDCs and their migration distance was significantly greater across Channel 4 and into Channel 5 in the FRC hydrogel relative to the blank hydrogel (Fig. 3A,C,D). The migrating TNF-moDCs in Channel 4 and in Channel 5 of the FRC hydrogel at day 3 were shown to be distributed throughout the majority of the Z-planes of the device (Fig. S6, Fig. S7). While the majority of the TNF-moDCs were not in contact with the FRCs (Fig. S6, Fig. S7), a proportion of the TNF-moDCs were shown to be in contact with the FRCs in Channel 4 and in Channel 5 at day 3 (Fig. 3B). Given its high expression by the FRCs (Fig. 2D, Fig. S4) and its reported role in DC migration (*17*), we explored the role of CCL2 on TNF-moDC driven migration into the FRC hydrogel with an antagonist against the CCL2 receptor (CCR2). The addition of a CCR2 antagonist to the FRC hydrogel led to no significant differences in TNF-moDC migration across Channel 4 and into Channel 5 relative to a FRC hydrogel with DMSO (Fig. 3E).

**Fig. 3.**
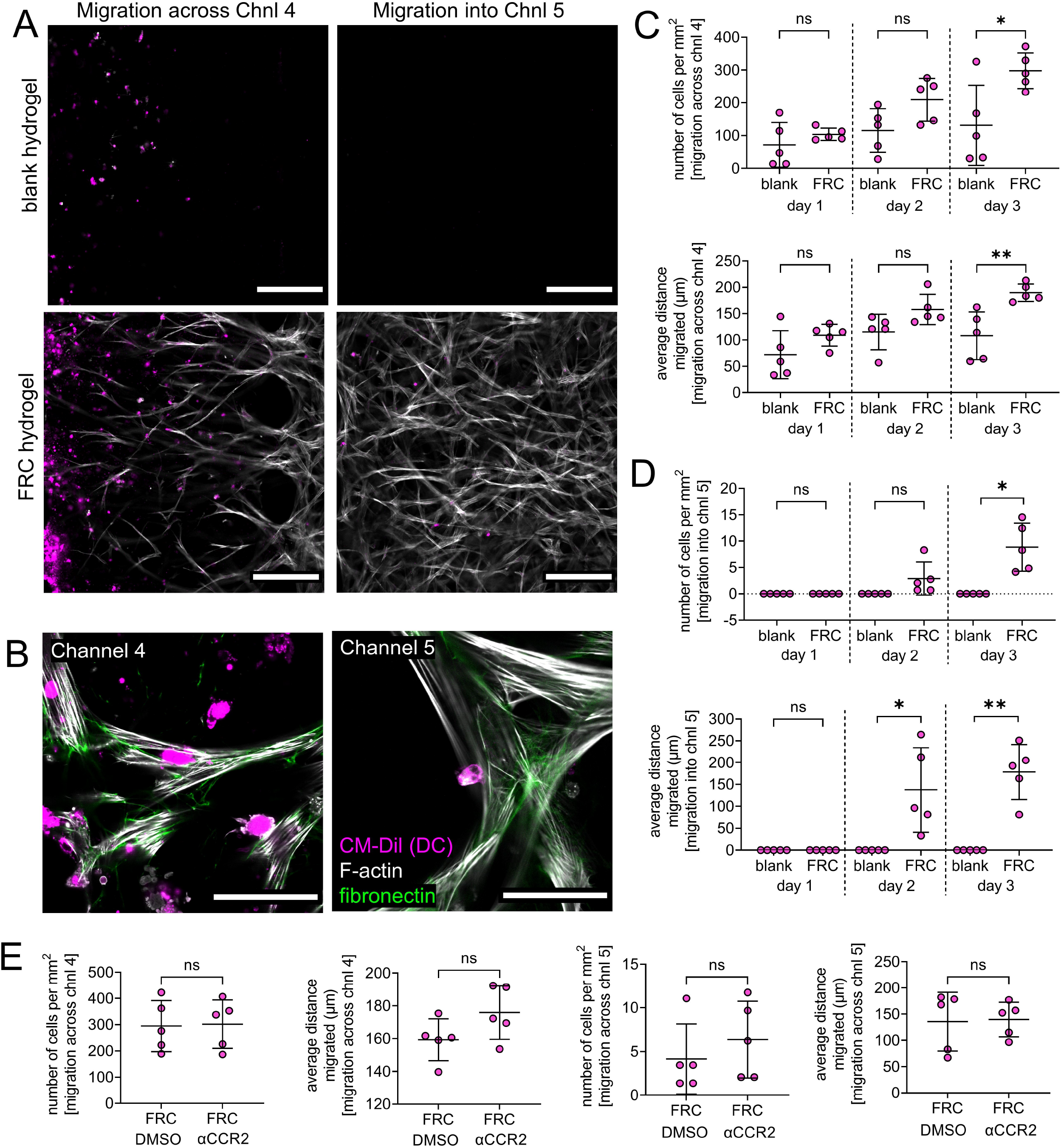
TNF-α-matured monocyte-derived dendritic cell migration toward lymph node stromal network. (A) Representative image of TNF-α-matured monocyte-derived dendritic cells (TNF-moDCs) migrating across Channel 4 and into Channel 5 toward a blank hydrogel and an FRC hydrogel at day 3. Scale bars = 200 μm. (B) Representative image of TNF-moDCs in Channel 4 and Chanel 5 in contact with FRCs at day 3. Magenta = CM-Dil labeled TNF-moDCs, white = phalloidin F-actin, green = fibronectin. Scale bar = 50 μm. (C,D) Quantification of TNF-moDC migration (C) across Channel 4 and (D) into Channel 5 over 3 days. n = 5 devices. (E) Quantification of TNF-moDCs migration across Channel 4 and into Channel 5 after 3 days into an FRC hydrogel with and without a CCR2 antagonist. n = 5 devices. Data represented as mean ± SD. Significance is denoted by \**P* < 0.05 or \*\**P* < 0.01 by with a two-tailed Student’s *t* test with or without Welch’s correction where applicable.

### Migration of T-cells toward lymph node stromal network

The migration of pan CD3^+^ human T-cells toward the lymph node stromal network was similarly evaluated by encapsulating the T-cells in a hydrogel in Channel 3 (Fig. 1B,C) and comparing their migration across Channel 4 and into Channel 5 (Fig. S5) over 3 days in a blank hydrogel versus a FRC hydrogel. At day 1, the average distance migrated across Channel 4 and into Channel 5, as well as the number of pan T-cells migrating into Channel 5 were significantly greater in the FRC hydrogel relative to the blank hydrogel (Fig. 4C,D). By day 2, similar observations were made, as the number of pan T-cells migrating across Channel 4 and into Channel 5, as well as the average distance migrated into Channel 5 was significantly greater in the FRC hydrogel relative to the blank hydrogel (Fig. 4C,D). At day 3, the average distance migrated across Channel 4 and the number of pan T-cells migrated into Channel 5 was significantly greater in the FRC hydrogel relative to the blank hydrogel (Fig. 4A,C,D). Migrating pan T-cells at day 3 in the FRC hydrogel were shown to be present in all Z-planes of Channel 4 and Channel 5 (Fig. S8, Fig. S9). A fraction of these migrating pan T-cells at day 3 were shown to be in physical contact with the FRCs (Fig. 4B).

**Fig. 4.**
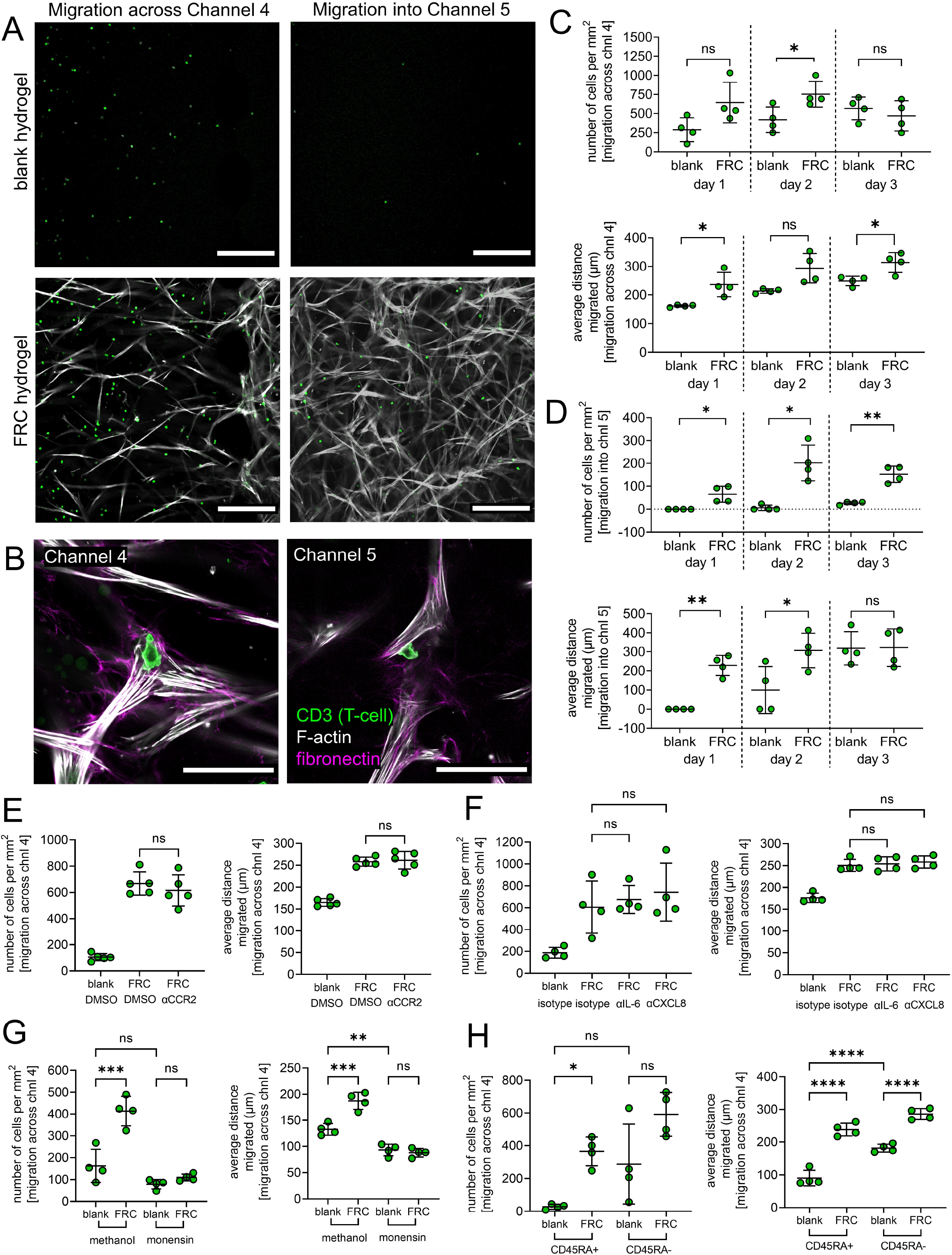
T-cell migration toward lymph node stromal network. (A) Representative image of pan T-cells migrating across Channel 4 and into Channel 5 toward a blank hydrogel and a FRC hydrogel at day 3. Scale bar = 200 μm. (B) Representative image of pan T-cells in Channel 4 and in Chanel 5 in contact with FRCs at day 3. Green = CD3 labeled T-cells, white = phalloidin F-actin, magenta = fibronectin. Scale bar = 50 μm. (C,D) Quantification of pan T-cell migration across Channel 4 and (D) into Channel 5 over 3 days. n = 5 devices. (E) Quantification of pan T-cell migration across Channel 4 toward a blank hydrogel and a FRC hydrogel with or without a CCR2 antagonist after 1 day. n = 5 devices. (F) Quantification of pan T-cell migration across Channel 4 toward a blank hydrogel and a FRC hydrogel in the presence of an isotype control antibody, anti-IL-6 antibody, and anti-CXCL8 antibody after 1 day. n = 4 devices. (G) Quantification of pan T-cell migration across Channel 4 toward a blank hydrogel and a FRC hydrogel with or without monensin after overnight incubation. n = 4 devices. (H) Quantification of CD45RA^+^ or CD45RA^-^ T-cell migration across Channel 4 toward a blank hydrogel and a FRC hydrogel after 1 day. n = 4 devices. Data are presented as means ± SD. For the data in (C,D), significance is denoted by \**P* < 0.05 or \*\**P* < 0.01 by with a two-tailed Student’s *t* test with or without Welch’s correction where applicable. For the data in (E-H), significance is denoted by \**P* < 0.05, \*\**P* < 0.01, \*\*\**P* < 0.001 or \*\*\**P* < 0.001 by one-way analysis of variance (ANOVA) with Tukey’s post hoc test.

Given the high concentrations of CXCL-8, IL-6, and CCL2 in the conditioned media of the FRC hydrogel network (Fig. 2D, Fig. S4) and their reported role in T-cell chemotaxis (*20-22*), we then evaluated the functional role of these chemokines and cytokines in mediating pan T-cell migration in the lymph node stromal network. Addition of a CCR2 antagonist to the FRC hydrogel did not significantly affect the migration of pan T-cells across Channel 4 and into Channel 5 relative to an FRC hydrogel with DMSO after 1 day (Fig. 4E, Fig. S10A). Similarly, addition of an IL-6 or a CXCL-8 neutralizing antibody to the FRC hydrogel did not significantly affect the migration of the pan T-cells across Channel 4 and into Channel 5 relative to an FRC hydrogel with an isotype control antibody after 1 day (Fig. 4F, Fig. S10B). We then tested the effect of adding monensin to pan T-cell migration, a protein transport inhibitor which decreases extracellular protein secretion from cells (*27*). With the addition of monensin, the migration of pan T-cells across Channel 4 and into Channel 5 was not significantly different between the FRC hydrogel and the blank hydrogel (Fig. 4G, Fig. S10C). In contrast, the average migration distance across Channel 4 and the number of pan T-cells migrating across Channel 4 and into Channel 5 was significantly greater in the FRC hydrogel relative to the blank hydrogel in the presence of the methanol control (Fig. 4G, Fig. S10C). The presence of monensin moderately influenced the spontaneous migration of the pan T-cells, as the average distance migrated across Channel 4 by the pan T-cells was significantly decreased in the blank hydrogel with monensin relative to the blank hydrogel with control methanol (Fig. 4G).

The ability of CD45RA^+^ (naïve) T-cells and CD45RA^-^ (central and effector memory) T-cells to migrate toward the lymph node stromal network was also evaluated. The number of CD45RA^+^ T-cells migrating across Channel 4 and the average distance migrated across Channel 4 and into Channel 5 was significantly greater in the FRC hydrogel relative to the blank hydrogel after 1 day (Fig. 4H, Fig. S10D). Similarly, the number of CD45RA^-^ T-cells migrating across Channel 5 and the average distance migrated across Channel 4 was significantly greater in the FRC hydrogel relative to the blank hydrogel after 1 day (Fig. 4H, Fig. S10D). The CD45RA^-^ T-cells exhibited greater spontaneous migration relative to the CD45RA^+^ T-cells, as the average distance migrated across Channel 4 was significantly greater for the CD45RA^-^ T-cells in the blank hydrogel relative to the CD45^+^ T-cells in the blank hydrogel (Fig. 4H).

### Co-migration of dendritic cells and T-cells toward the lymph node stromal network

To evaluate co-recruitment of TNF-moDCs and T-cells toward the lymph node stromal network, we utilized a custom designed, polydimethylsiloxane microfluidic device that spatially isolates the initial seeding of TNF-moDCs, T-cells, and FRCs (Fig. 5A,B). The same protocol for forming the lymph node stromal network followed by addition of immune cells to the device was followed (Fig. 1C) and the migration of immune cells into Channel 5 in a blank hydrogel and FRC hydrogel was evaluated (Fig. S11). Co-recruitment of TNF-moDCs and pan T-cells isolated from a single donor was first evaluated. At day 3, the number of TNF-moDCs migrating into Channel 5 was significantly greater in the FRC hydrogel relative to the blank hydrogel (Fig. 5D); no significant differences were observed for the pan T-cells migrating into Channel 5 between the blank and the FRC hydrogel (Fig. 5D). The TNF-moDCs and pan T-cells were co-recruited to the same region of Channel 5 throughout the majority of the Z-planes of the device (Fig. 5C, Fig. S11). While physical contact was observed between the FRCs and the immune cells (Fig. 5C), no physical contact was observed between the TNF-moDCs and the pan T-cells.

**Fig. 5.**
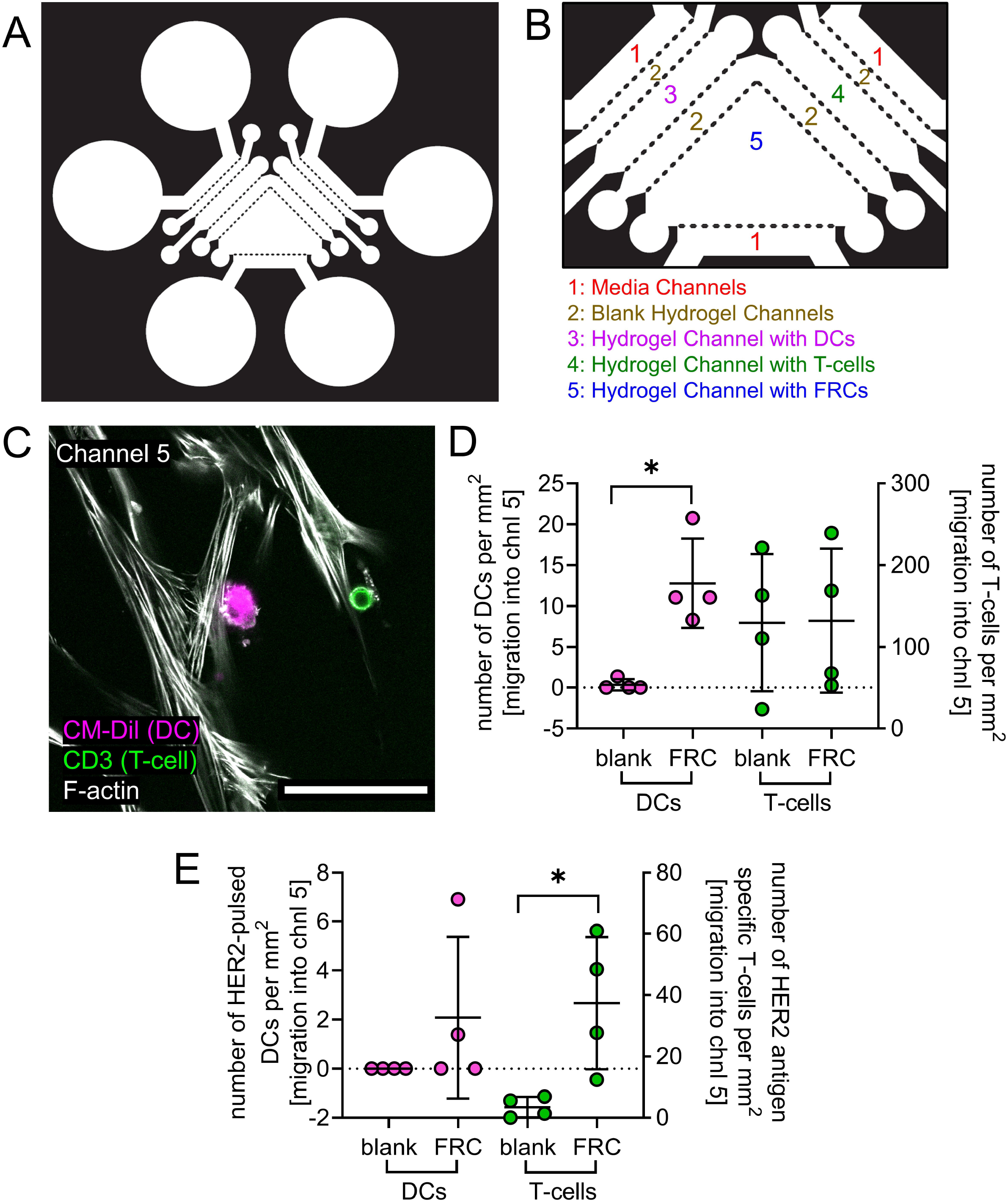
Dual migration of TNF-α-matured monocyte-derived dendritic cells (TNF-moDC) and T-cells toward lymph node stromal network. (A) Photolithography mask of device for fabricating the tri-culture lymph node stromal network chip. (B) Zoomed in image of photolithography mask of tri-culture device, denoting channel numbers and channel contents. (C) Representative image of a TNF-moDC and pan T-cell in Channel 5. Magenta = CM-dil labeled T-DCs, Green = CD3 labeled T-cells, white = phalloidin F-actin. Scale bar = 50 μm. (D) Quantification of TNF-moDC and pan T-cell migration into Channel 5 in a blank hydrogel and FRC hydrogel at day 3. n = 4 devices. (E) Quantification of TNF-moDC pulsed with HER2/neu (369-377) and CD8^+^ HER2/neu antigen-specific T-cell migration into Channel 5 in a blank hydrogel and FRC hydrogel at day 3. n = 4 devices. Significance is denoted by \**P* < 0.05 by with a two-tailed Student’s *t* test with or without Welch’s correction where applicable.

The ability of the lymph node stromal network to co-recruit TNF-moDCs pulsed with HER2/neu peptide and CD8^+^ HER2/neu specific T-cells was also evaluated. At day 3, the number of CD8^+^ HER2/neu specific T-cells recruited to Channel 5 was significant greater in the FRC hydrogel relative to the blank hydrogel (Fig. 5E). There were no significant differences in the recruitment of TNF-moDCs pulsed with HER2/neu (369-377) peptide between the FRC hydrogel and the blank hydrogel (Fig. 5E). No physical contacts were observed between TNF-moDCs and the HER2/neu specific T-cells.

## Discussion

While there are extensive studies evaluating the function of FRCs *in vivo* in murine models and *in vitro* on TCPS, there is a need for 3D models of the human paracortex that can evaluate the trafficking of human DCs and T-cells into the lymph node. Although many molecular pathways have been shown to be conserved between human and mouse FRCs, human FRCs have been shown to differentially express a variety of cytokines and inflammatory pathways compared to mouse FRCs (*13, 28*). Thus, in comparison to *in vivo* mouse models, *in vitro* models of the lymph node paracortex utilizing human FRCs may serve as better predictive models for preclinically testing immunotherapies targeting lymph nodes. Here, we engineered an on-chip lymph node stromal network from human FRCs in multi-channel microfluidic devices to enable co-culture of FRCs and lymphocytes. The engineered lymph node stromal network, consisting of an interconnected network of FRCs, secreted extracellular matrix proteins and an array of chemokines implicated in lymphocyte trafficking. TNFα-moDCs, CD45RA^+^ T-cells, and CD45RA^-^ T-cells migrated toward the lymph node network to a greater extent than toward a blank hydrogel, likely due to the combined effects of multiple cytokines secreted by the FRCs. Both TNFα-moDCs and pan T-cells physically interacted with the FRC network. Furthermore, TNFα-moDC and T-cell (both pan-T cells and antigen-specific T-cells) co-recruitment to the FRC network was demonstrated in the tri-culture device.

In our microfluidic collagen I-fibrin composite hydrogel, the primary, human CD31^-^ CD45^-^PDPN^+^ FRCs formed an interconnected cell network, as well as deposited entactin, collagen IV, and fibronectin into the surrounding hydrogel. Previous work has demonstrated that fibronectin and collagen IV are expressed on the outer membrane surface of FRCs in the lymph nodes of cynomolgus monkeys (*29*). Entactin, collagen IV, and fibronectin have also been observed in the reticular fibers of human lymph nodes (*30*). The FRC network formed in our microfluidic system, however, exhibits several distinct differences from *in vivo* lymph node paracortex in terms of geometry and architecture. In the lymph node paracortex of mice, the spacing between FRC strands was shown to be between 5 μm to 37 μm (average of 17 μm) (*6*), which is smaller than the average spacing between the FRCs in our engineered system (Fig. S2B). Furthermore, FRCs in the lymph node paracortex of mice, humans, and cynomolgus monkeys ensheath an extracellular matrix network made of various proteins, including collagen III and laminin, that also serves as a conduit system for lymph molecules (*2, 29, 30*). In our model, neither collagen III, laminin, nor a distinct conduit network were secreted by the FRCs.

Future optimization of culture conditions and biomaterial hydrogel composition may be necessary to recapitulate these other structural features of the *in vivo* lymph node paracortex.

The self-assembled FRC network was shown to actively secrete a variety of chemokines and cytokines, including CCL2, IL-6, CXCL8, and CXCL1, several of which have been shown to regulate dendritic cell and T-cell trafficking. Previous work has demonstrated that FRCs in the lymph node paracortex of mice similarly express CCL2, which regulates monocyte trafficking to the paracortex (*31*). Furthermore, human cultured FRCs derived from lymph nodes or tonsils have also been shown to constitutively express CCL2, IL-6, CXCL8, and/or CXCL1 (*15, 16, 28*). Notably, FRCs in the engineered stromal network did not express characteristic CCL19, CCL21, CXCR4, or IL-7, which have been shown to induce the recruitment of mature dendritic cells and naïve T-cells to the lymph node paracortex and induce naïve T-cell survival (*5, 32*). CCL19 and CCL21 expression has commonly been shown to be downregulated or lost in cultured human FRCs (*13, 15, 16*). CCL21 expression in primary mouse FRCs, however, has been shown to be maintained by culturing the FRCs under flow conditions (*10*). Furthermore, FRC subsets from the mouse lymph nodes have been shown to exhibit heterogeneity in their secretory factors (*33*). Utilizing freshly isolated human FRCs in our microfluidic device with interstitial flow may reconstitute some of these missing cytokines in our system.

Despite the lack of CCL19 or CCL21 expression by the FRCs in our system, TNFα-moDCs migrated toward the lymph node stromal network to a greater extent than toward a blank hydrogel, where a portion of the TNFα-moDCs were in contact with the FRC network. This enhanced migration toward the lymph node stromal network was not shown to be dependent on the sole activity of CCR2, a receptor for CCL2, on the TNFα-moDCs. Immature monocyte derived dendritic cells have been shown to actively migrate toward human lymph node FRCs via the activity of CCL2 and CCL20 secreted by FRCs (*15*). While migration of the TNFα-moDCs did not show the same dependency on CCL2 for their migration, we postulate that the enhanced migration of the TNFα-moDCs toward the FRC network may be due to the combined action of several chemokines secreted by the FRCs. These cytokines include GM-CSF, CCL5, CCL7, and CCL3, which have previously been shown to drive dendritic trafficking (*18, 19*). The physical contacts between the TNFα-moDCs and the FRC network may be due to the ability of DCs to adhere or interact with the surface of FRCs via distinct molecular cues. In mice, murine DCs were shown to migrate along FRCs via the interaction between CLEC-2 on the DCs with PDPN on the FRCs (*34*). Furthermore, mature, murine dendritic cells were shown to migrate toward and adhere to FRC cell lines derived from BALB/c mice *in vitro* (*9*).

Human pan T-cells, including CD45RA^+^ naïve T-cells cells and CD45RA^-^ effector/memory T-cells, migrated toward the lymph node stromal network to a greater extent than toward a blank hydrogel; the pan T-cells were shown to be in contact with the FRCs. This directed migration was shown to not depend on the action of CCR2 ligands, CXCL8, or IL-6 alone, but likely on the combined action of multiple cytokines secreted from the FRCs based on our monensin experiments. Other T-cell migratory cytokines expressed by the FRC network include CCL5 (*23*), CCL7 (*24*), CCL3 (*25*), and CXCL10 (*26*). Intravital microscopy of mouse lymph nodes has shown that T-cells actively migrate along FRC networks of the lymph node paracortex (*6*). Furthermore, it has long been appreciated that naïve T-cells and subsets of memory T-cells home to lymph nodes (*35, 36*). Furthermore, murine CD4^+^ T-cells actively migrated toward and adhered to FRC cell lines derived from BALB/c mice *in vitro* (*9*). The adhesion of the pan T-cells to the FRC network may be mediated by distinct molecular cues, such as VCAM-1 expression on FRCs (*37*).

In our tri-culture system, co-recruitment of TNFα-moDCs and pan or antigen-specific T-cells to the FRC network was enhanced relative to a blank hydrogel. However, antigen presentation of DCs to T-cells or activation of antigen-specific T-cells was not readily observed. Previous work, however, has demonstrated the importance of FRCs to T-cell activation by DCs. Depletion of FRCs before the initiation of an influenza virus infection was shown to reduce naïve T-cell trafficking, dendritic cell recruitment, and T-cell activation in mice (*38*). The lack of observable antigen-presentation in our model may be due to the longer periods of time the TNFα-moDCs migrated into the FRC channel, which was approximate 2-3 days. This may have resulted in their reduced activation over time. Furthermore, the number of TNFα-moDCs and T-cells reaching the FRC channel may have been too low to observe stochastic interactions and activation between the DCs and T-cells.

The results of this study have yielded an engineered stromal network that can evaluate the ability of human TNFα-moDCs and T-cells to migrate toward an engineered human lymph node paracortex. Future work will explore how to more accurately recapitulate the structure and function of the human lymph node paracortex, in terms of the ability of the FRCs to secrete key chemokines, such as CXCR4, IL-7, and CCR7-ligands, and produce a conduit system. These improvements and optimizations may long term yield a system that can simultaneously evaluate DC and T-cell migration and antigen presentation. Furthermore, this model may serve as a basis to engineer assays that can evaluate the potency of human DC and T-cell immunotherapies, specifically in terms of their ability to migrate toward the lymph node paracortex via CCR7-independent mechanisms.

## Methods

### Culture and wxpansion of juman fibroblastic reticular cells

Passage 0 human lymphatic fibroblasts (ScienCell, Cat. # 2530) were cultured in complete FRC media (α-MEM media [Gibco, Cat. # 12571] supplemented with 16% FBS, 1% P/S, and L-GlutaMAX™) at an initial seeding density of 1,430 cells/mm^2^. Upon reaching 80-90% confluency, the fibroblasts were passaged with TrypLE™ Express Enzyme (ThermoFisher Scientific Cat. # 12604013) and stained with the following antibodies in FACS staining buffer (PBS with 2% Fetal Bovine Serum, 1% Penicillin-Streptomycin): anti-human CD31 (BioLegend, WM59, FITC), anti-human CD45 (BioLegend, 2D1, PE), and anti-human podoplanin (BioLegend, NC-08, APC). Human lymphatic fibroblasts were then enriched by FACS for CD31^-^CD45^-^podoplanin^+^ fibroblastic reticular cells (Fig. S1B) into FACS sorting buffer (PBS with 0.5% bovine serum albumin, 2 mM EDTA, and 1% Penicillin-Streptomycin) using a BD FACSAria™ Fusion Cell Sorter. The sorted fibroblastic reticular cells were then expanded in complete FRC media, passaged, frozen down, and further expanded at an initial seeding density of 1,430 cells/mm^2^. Upon reaching 80-90% confluency, the fibroblastic reticular cells were passaged prior to incorporation into the microfluidic devices.

### Generation of TNF-α matured monocyte-derived dendritic cells

CD14 microbeads (Miltenyi Biotec, Cat. # 130-050-201) were used to isolate CD14^+^ cells from peripheral blood mononuclear cells (PBMCs) [All Cells, Cat.# LP, FR, MNC, 300M)]. CD14^+^ cells were then cultured for 5-6 days in RPMI1640 (ATCC, Cat.# 30-201) supplemented with 10% FBS and 1% P/S at 5×10^5^ cells/mL with 800 IU/mL GM-CSF (R&D, Cat. # 215-GM-050) and 500 IU/mL IL-4 (R&D, Cat. # 204-IL-020), with fresh media and cytokines added on day 2 and day 4. On day 5-6, immature dendritic cells were matured in the presence of 20 ng/mL TNF-α (R&D, Cat. #), 800 IU/mL GM-CSF, and 500 IU/mL IL-4 for 48 hours. In experiments with antigen specific T-cells, the dendritic cells were pulsed with 10 µg/mL HER2/neu (369-377) peptide (Cellero, Cat. # 1136) overnight prior to collection of the cells. Cells were then collected and stained with Vybrant™ CM-Dil Cell-Labeling Solution (ThermoFisher Scientific, Cat. # V22888) according to manufacturer’s instructions. In experiments with antigen specific T-cells, the dendritic cells were pulsed again with 100 µg/mL HER2/neu (369-377) peptide (Cellero, Cat. # 1136) for one hour at 37ºC.

### Pan, naïve, and anti-specific T-cells

Primary pan T-cells were isolated from PBMCs with the Pan T-Cell Isolation Kit, human (Miltenyi Biotec, Cat. # 130-096-535) according to manufacturer’s instructions. Naïve, CD45RA^+^ T-cells were isolated from CD3^+^ enriched pan T-cells with the Naïve Pan T Cell Isolation Kit, human (Miltenyi Biotec, Cat. # 130-097-095) according to manufacturer’s instructions; the remaining cells from the CD3^+^ enriched pan T-cells were considered to be CD45RA^-^ T-cells. CD8^+^ anti-HER2/neu (369-377) antigen-specific T-cells were obtained commercially from Charles River (Charles River, Cat. # ASTC-1126).

### Microfluidic device fabrication

Microfluidic devices made of polydimethylsiloxane (PDMS) were fabricated with soft lithography. SU-8 100 negative photoresist (Kayaku Advanced Materials, Cat. # Y131273) was spun to generate a 125 μm thick layer on a silicon wafer. The channel designs, printed by Fineline Imaging Inc., were cured onto the SU-8 layer with UV light on an OAI Model 200 Mask Aligner to form the channel layer. Dow Sylgard™ 184 Silicone Encapsulant Clear (Ellsworth Adhesives, Cat. # 4019862) was mixed using a curing ratio of 10:1, poured over the master mold, and cured on an 80°C hotplate for 3 hours. The polymerized PDMS was then peeled off the mold and cut into individual devices. For the co-culture devices (Fig. 1A), an 8 mm and a 1 mm biopsy punch were used to punch out the media reservoirs and hydrogel ports respectively. For the tri-culture devices (Fig. 5A), a 6 mm and a 1 mm biopsy punch were used to punch out the media reservoirs and hydrogel ports respectively. The PDMS devices were then bonded to a glass cover slip by plasma treatment and were kept in an 80°C dry oven overnight to restore hydrophobicity. UV radiation was used to sterilize devices prior to addition of hydrogels.

### 3D lymph node chip hydrogel and culture

A collagen I-fibrin composite hydrogel was used as a 3D scaffold to engineer the lymph node stromal chip. To fabricate the hydrogel, a 4 mg/mL collagen I gel was first prepared by mixing type I bovine atelocollagen (Advanced Biomatrix, Cat. # 5133), 10X PBS (1/10 final volume), 1 N sodium hydroxide (0.0023 × volume of collagen added), and distilled water on ice. The resulting collagen solution was then mixed in a 1:1 ratio with 1 U/mL bovine thrombin solution (Sigma, Cat. # T4648) in PBS. The resulting collagen I-thrombin solution was then mixed in a 1:1 ratio with 3 mg/mL bovine fibrinogen (Sigma, Cat. # F8630) solution in PBS, yielding a hydrogel with a final concentration of 1.5 mg/mL fibrinogen, 0.25 U/mL thrombin, and 1 mg/mL collagen I. The resulting hydrogel was cross-linked for 1 hour at 37°C.

To fabricate the co-culture device (Fig. 1A) and tri-culture device (Fig. 5A), a blank collagen-fibrin hydrogel was first injected into Channels 2 and 4 of the co-culture device and into all Channel 2 lanes in the tri-culture device. After cross-linking the resulting hydrogel for 1 hour at 37°C, a collagen-fibrin hydrogel embedded with 15×10^6^ FRCs/mL was then injected into Channel 5 of both devices. After cross-linking the device for another 1 hour at 37°C, complete FRC media was added to all remaining channels in the devices. After 4 days of culture, media from all blank channels was removed and a collagen I-fibrin hydrogel with 8 million DCs/mL was injected into Channel 3 of the co-culture device. In other experiments, a collagen I-fibrin hydrogel with 20 million T-cells/mL was injected in Channel 3 of the co-culture device. In the tri-culture device, a collagen I-fibrin hydrogel with 8 million DCs/mL was added in Channel 3 and a collagen I-fibrin hydrogel with 20 million T-cells/mL was injected in Channel 4. In all experiments with DCs, T-cells, both cell types, or no immune cells, the media in channels was switched to immune cell media (RPMI1640 media supplemented with 10% FBS and 1% P/S) after 4 days of initial culture and were then cultured for up to 3 additional days.

### Cytokine Array

After culturing the collagen I-fibrin microfluidic device for four days, the media in the culture with FRCs and a blank hydrogel was switched to RPMI1640 media supplemented with 1% FBS and 1% P/S. Media was collected and changed with fresh media every 24 hours for 3 days. Cytokine and chemokine concentrations in the conditioned media were analyzed with the Human Cytokine/Chemokine 71-Plex Discovery Assay® Array (Eve Technologies, Cat. # HD71) and were multiplied by the total media volume to convert cytokine concentrations to picograms (pg) secreted per 24 hours. The concentration of cytokines and chemokines secreted by the FRCs were normalized by subtracting the average concentration of cytokines and chemokines detected in the conditioned media of blank hydrogel devices.

### Cytokine and chemokine inhibition experiments

Prior to adding immune cells to the lymph node chip after four days of culture, the media on the chip was switched to RPMI1640 media supplemented with 10% FBS and 1% P/S then incubated with the following antibodies or small molecules for at least 1 hour: 1 μM BMS CCR2 22 (R&D, Cat. # 3129, DMSO solvent), 5 μg/mL mouse IgG_1_ isotype control (R&D, Cat. # MAB002), 5 μg/mL anti-IL-6 (R&D, Cat. # MAB206, Mouse IgG_1_), 5 μg/mL anti-CXCL8 (R&D, Cat. # MAB208, Mouse IgG_1_), or 2 μM monensin (BioLegend, Cat. # 420701). After addition of DCs or T-cells to the device, fresh media with fresh antibodies and small molecules were added to the device for varying amounts of time.

### Immunohistochemistry and imaging

Lymph node stromal chips were fixed with 2% paraformaldehyde for 1 hour and then subsequently stained by immunohistochemistry. Nuclei were stained with TO-PRO™-3 Iodide (ThermoFisher Cat. # T3605) and F-actin with AlexaFluor™ 488 Phalloidin (ThermoFisher Cat. # A12379) or AlexaFluor™ 647 Phalloidin (ThermoFisher Cat. # A22287) according to manufacturer’s instructions. T-cells were fluorescently labeled with AlexaFluor 647 anti-human CD3 Antibody (BioLegend #300416) at a concentration of 5 μg/mL. The following extracellular matrix proteins were stained with the following primary antibodies: anti-fibronectin (Abcam, Cat. # ab2413), anti-vitronectin (Abcam, Cat. # ab45139), anti-tenascin C (Abcam, Cat. # ab108930), anti-collagen III (Abcam, Cat. # 184993), anti-collagen VI (Abcam, Cat. # ab182744), anti-entactin (Abcam, Cat. # ab254325), anti-laminin (Abcam, Cat. # ab11575), or anti-collagen IV (Abcam, Cat. # ab6586). Primary antibodies were stained with Goat anti-Rabbit IgG Secondary Antibody, AlexaFluor™ 488 (ThermoFisher, Cat. # A-11008), Goat anti-Rabbit IgG Secondary Antibody, AlexaFluor™ 568 (ThermoFisher, Cat. # A-11036), or Goat anti-Rabbit IgG Secondary Antibody, AlexaFluor™ 647 (ThermoFisher, Cat. # A27040) according to manufacturer’s instructions.

Images were taken on a Zeiss 710 Confocal Microscope under 10×, 20×, and 63× magnification. To quantify the number of cells per mm^2^ and the average cell distance migrated (μm) across Channel 4 and into Channel 5, 10× magnification Z-stack images (every 20 μm) were taken at regions of interest in Channel 4 and Channel 5 (Fig. S5, S11). Images quantifying “Migration across Channel 4” included a small portion of Channel 5 based on the size of the images. The Z-stack images were converted into maximum projection intensity images on Zeiss Zen Black software and the number of cells in the merged image were counted manually on FIJI (ImageJ) with the Cell Counter App and divided by the area of the image. Cell migration distances were quantified by converting the X-coordinates of the counted cells from pixels to μm.

### Statistics

Data presented in this study are expressed as means ± SD. Data were statistically analyzed in GraphPad Prism 9 (GraphPad Software Inc., CA). Significance between groups with two conditions was determined using unpaired, two-tailed Student’s *t* tests, with Welch’s correction when SDs were significantly different. Sample variance was tested using the *F* test. In groups with more than two conditions, significance between groups was determined using analyses of variance (ANOVAs) with Tukey’s post hoc test. Statistical significance was determined at *P* < 0.05.

## Acknowledgments

This work was supported in part by B.J.K.’s appointment to the Research Participation Program at CBER administered by the Oak Ridge Institute for Science and Education through the U.S. Department of Education and U.S. Food and Drug Administration. This work is also supported by B.J.K.’s appointment to the Interagency Oncology Task Force Fellowship through the National Cancer Institute and the U.S. Food and Drug Administration. This work was also partially supported by research funds from the Division of Cell Therapy 1.

## Author contributions

B.J.K. and K.E.S. conceived and designed the research. B.J.K. carried out the experiments. A.A. performed flow cytometry sorting. B.J.K. and K.E.S. wrote the paper. All authors reviewed and commented on the manuscript.

## Competing interests

The authors indicate no potential conflicts of interest.

## Corresponding Author

Correspondence to Kyung E. Sung.

## Data Availability

All study data are included in the article and/or supplementary information.

## Notes

### Competing Interest Statement

The authors have declared no competing interest.

