## Supplementary Information for "On-chip human lymph node stromal network for evaluating dendritic cell and T-cell trafficking"

|  | **Figures** | **Donor Information** | | |
| --- | --- | --- | --- | --- |
|  |  | **Gender** | **Age** | **Race** |
| Human Lymphatic Fibroblasts | All Figures | Female | 26 | N/A |
| PBMC derived MoDCs and T-cells | Fig. 3, 4, S7, S8, S9, S10, S11 | Female | 37 | Caucasian |
| PBMC derived MoDCs and T-cells | Fig. 5C, 5D, S12 | Male | 27 | Asian |
| MoDCs and HER2-specific T-cells | Fig. 5E | Male | 49 | Caucasian |

**Table S1.** List of all cell sources used with respective figure and donor information.

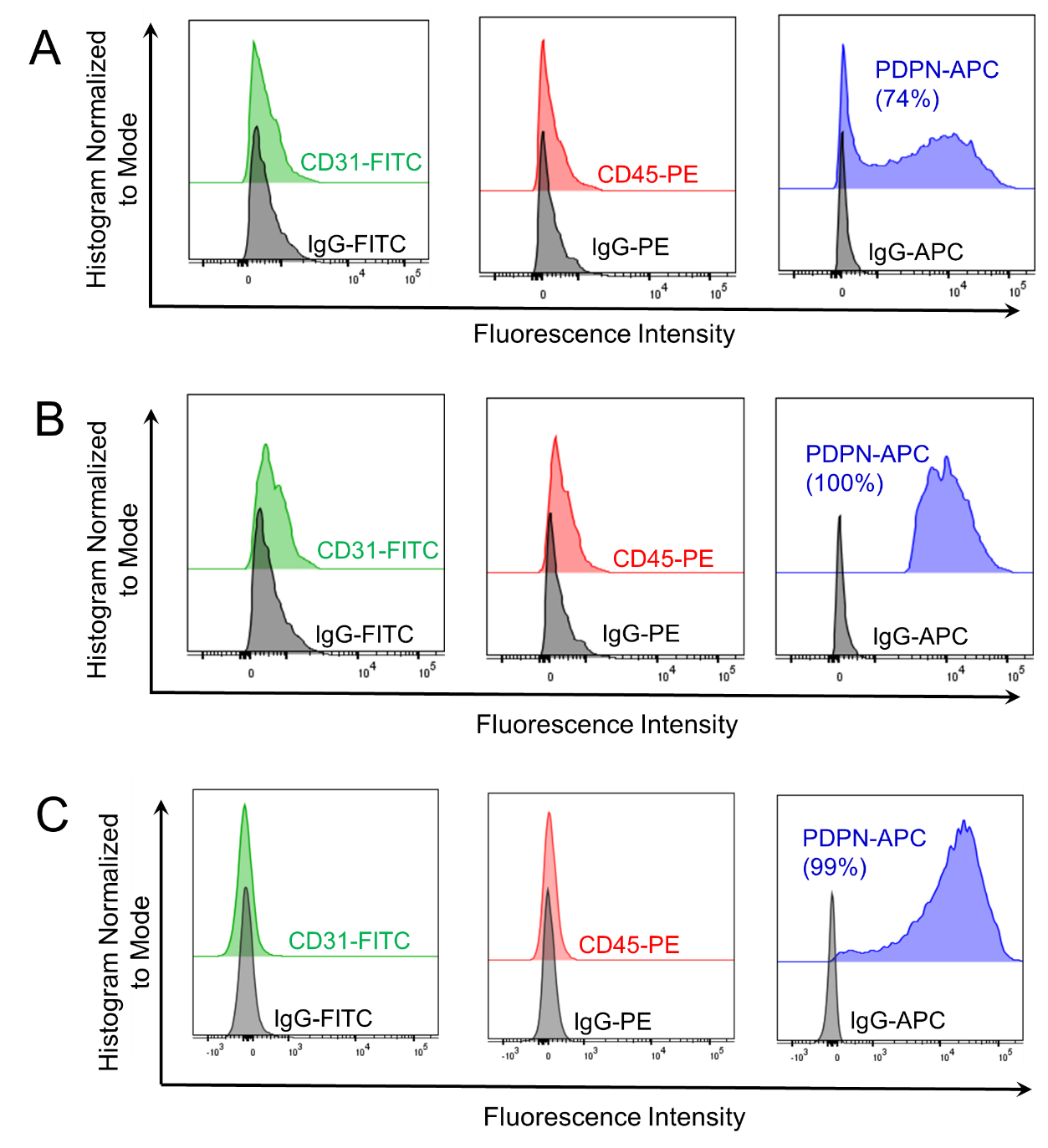

**Fig. S1.** **Flow cytometry analysis of human lymphatic fibroblasts.** (A) Flow cytometry histograms showing CD31, CD45, and podoplanin (PDPN) expression on human lymphatic fibroblasts prior to fluorescence activated cell sorting (FACS). (B) Flow cytometry histograms showing CD31, CD45, and PDPN expression on human lymphatic fibroblasts immediately after FACS enrichment of CD31^-^CD45^-^PDPN^+^ fibroblastic reticular cells. (C) Flow cytometry histograms showing CD31, CD45, and PDPN expression on fibroblastic reticular cells after expansion and immediately prior to incorporation into lymph node stromal chips.

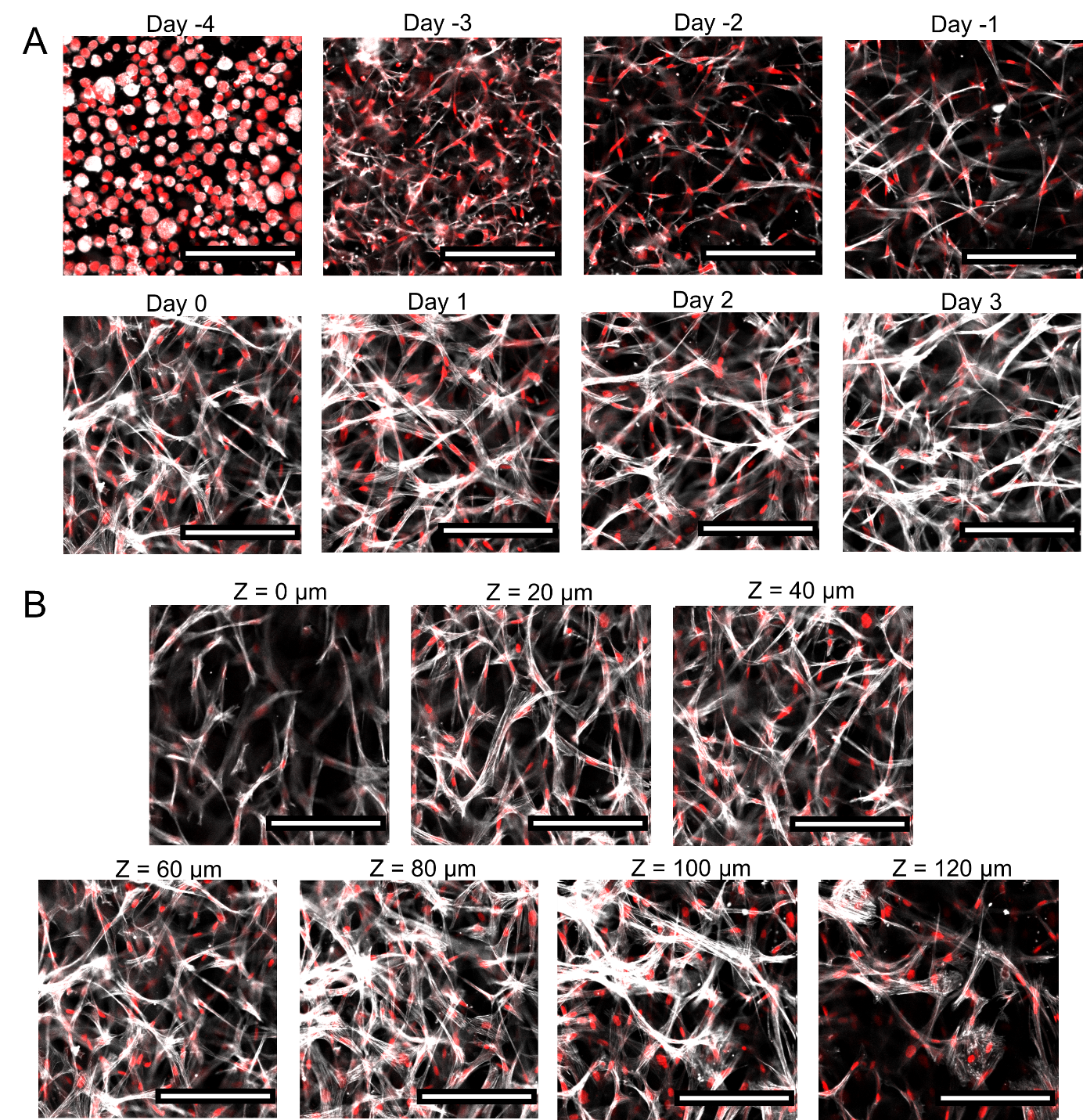

**Fig. S2. Fibroblastic reticular cell network formation in Channel 5.** (A) Representative confocal microscopy images of fibroblastic reticular cells spreading and forming an interconnected cell network in Channel 5 from day -4 to day 3. Media was changed from FRC media to immune cell media on day 0. (B) Confocal microscopy images of fibroblastic reticular cell network in Channel 5 on day 0 at different Z-stack planes. Red = nuclei, white = phalloidin F-actin. Scale bars = 200 μm.

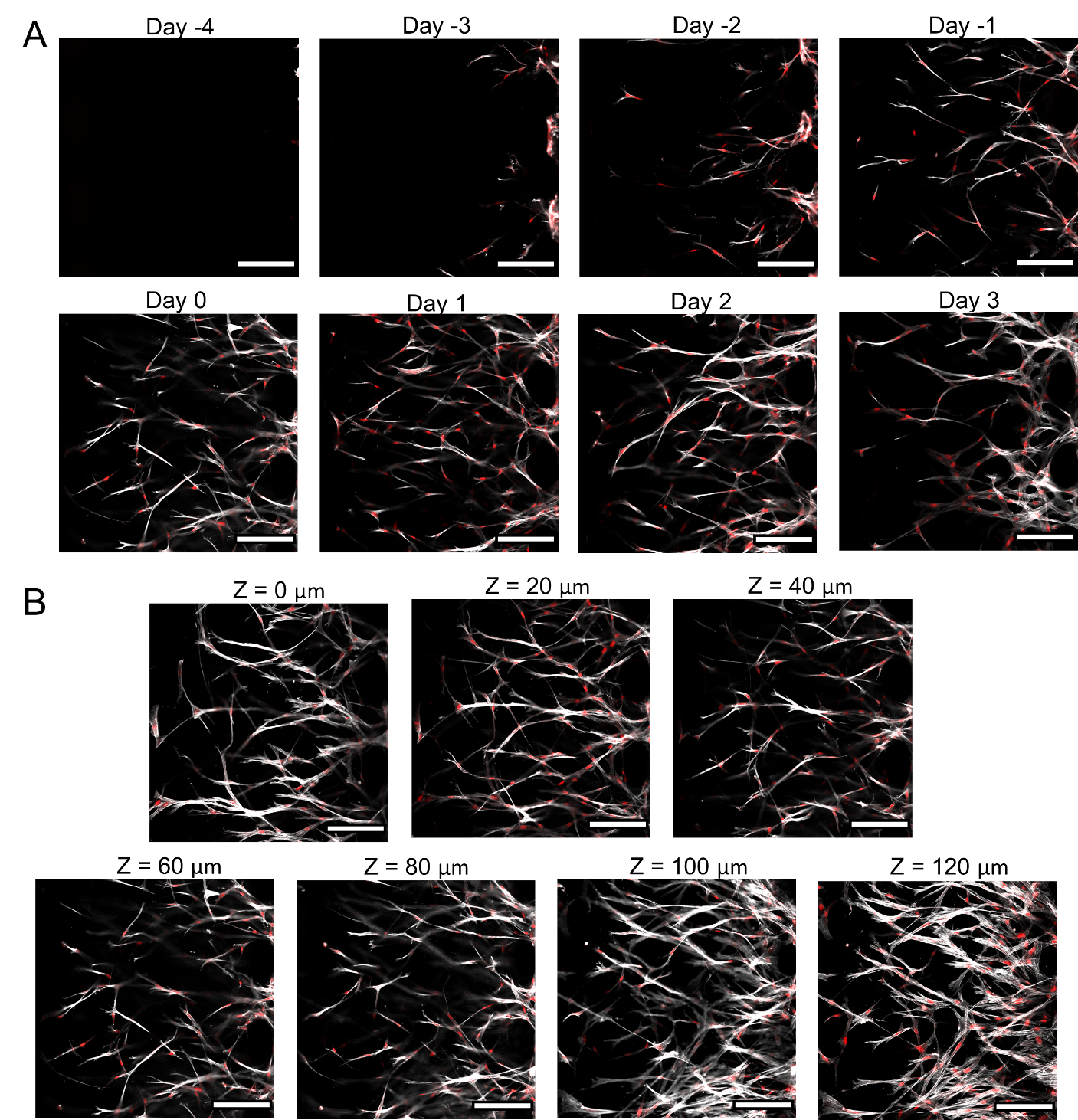

**Fig. S3. Fibroblastic reticular cell network formation in Channel 4.** (A) Representative confocal microscopy images of fibroblastic reticular cells spreading and migrating into Channel 4 from day -4 to day 3. Media was changed from FRC media to immune cell media on day 0. (B) Confocal microscopy images of fibroblastic reticular cells in Channel 4 on day 0 at different Z-stack planes. Red = nuclei, white = phalloidin F-actin. Scale bars = 200 μm.

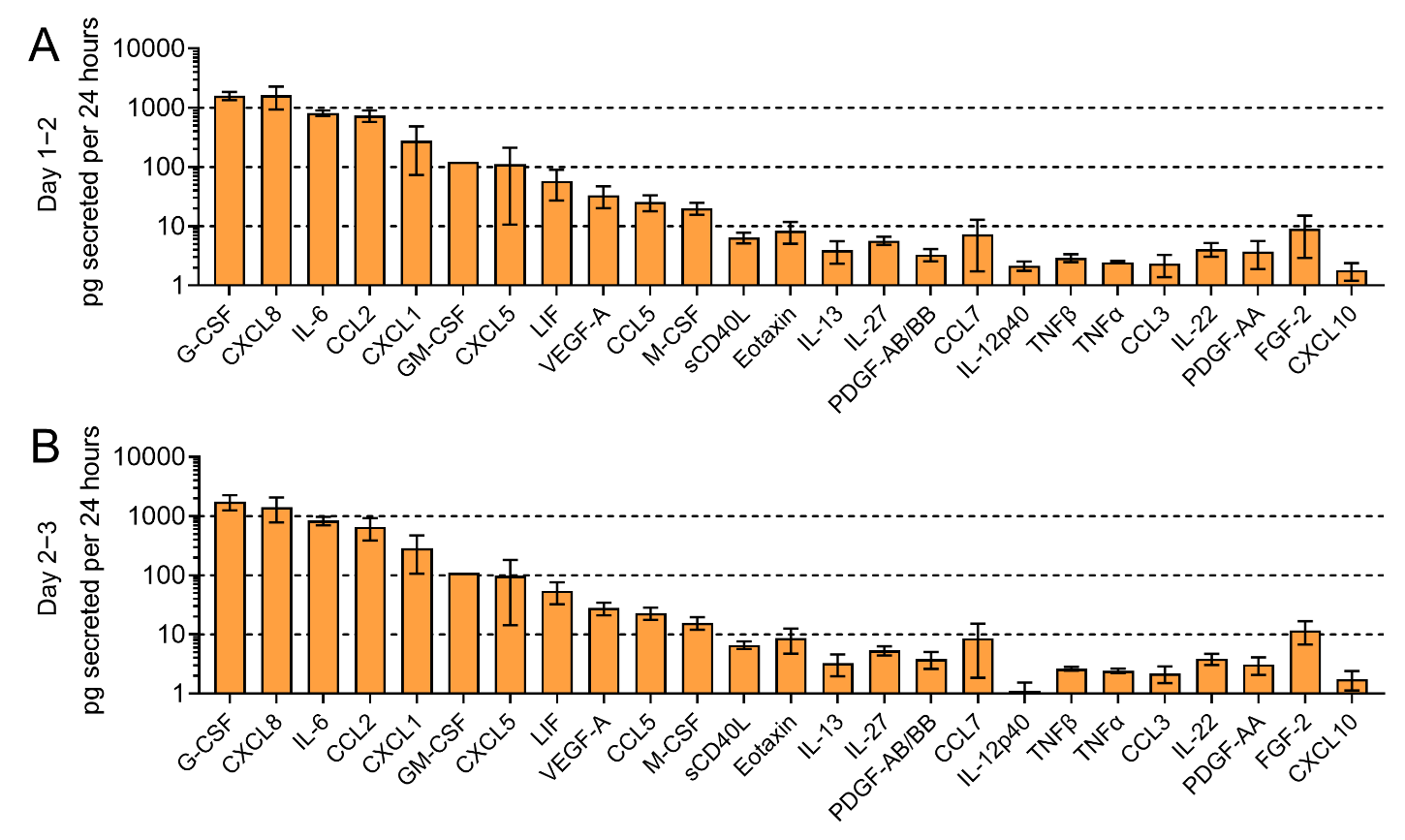

**Fig. S4. Secretome of fibroblastic reticular cells in lymph node paracortex device.** Picograms of chemokines and cytokines secreted by fibroblastic reticular cells in lymph node stromal chip over 24 hours from (A) day 1 to day 2 and (B) day 2 to day 3. n = 4 devices. Data are presented as means ± SD.

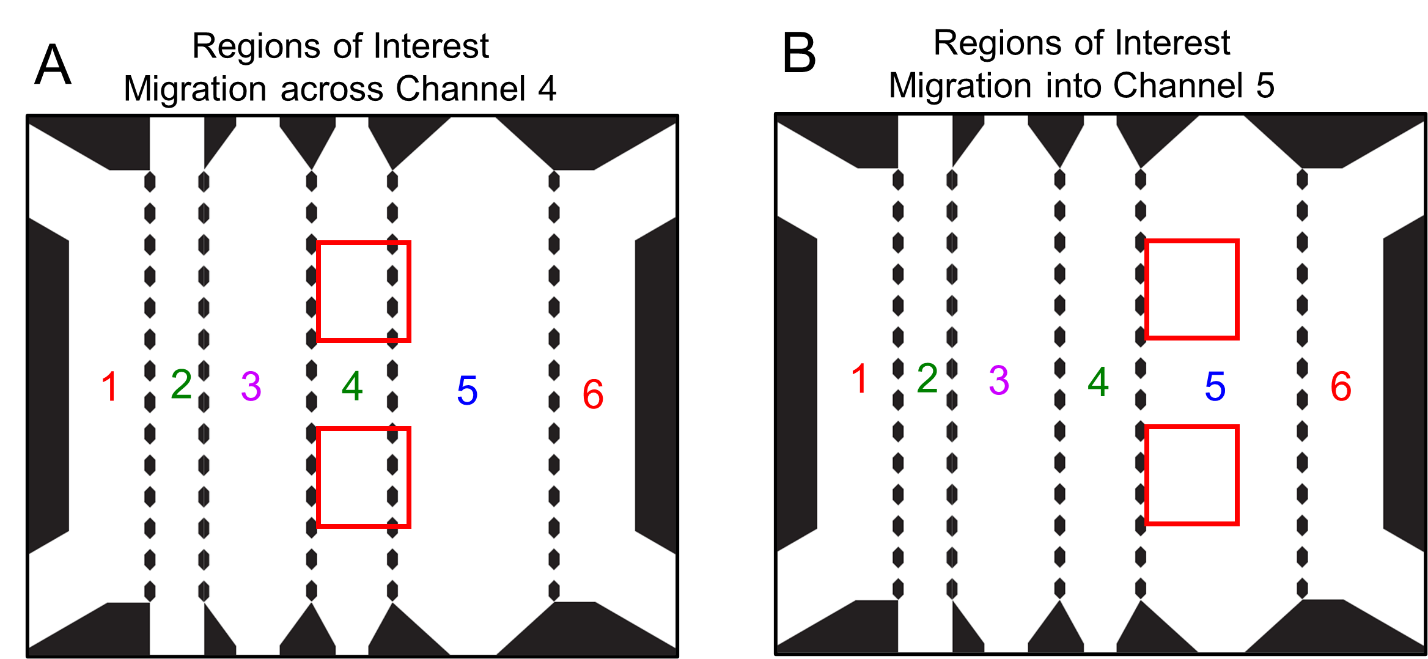

**Fig. S5. Regions of interest on lymph node stromal chip for quantifying dendritic cell and T-cell migration.** Zoomed in image of photolithography mask denoting regions in red boxes where confocal Z-stack images were taken across all devices and experiments to quantify (A) migration across Channel 4 and (B) migration into Channel 5.

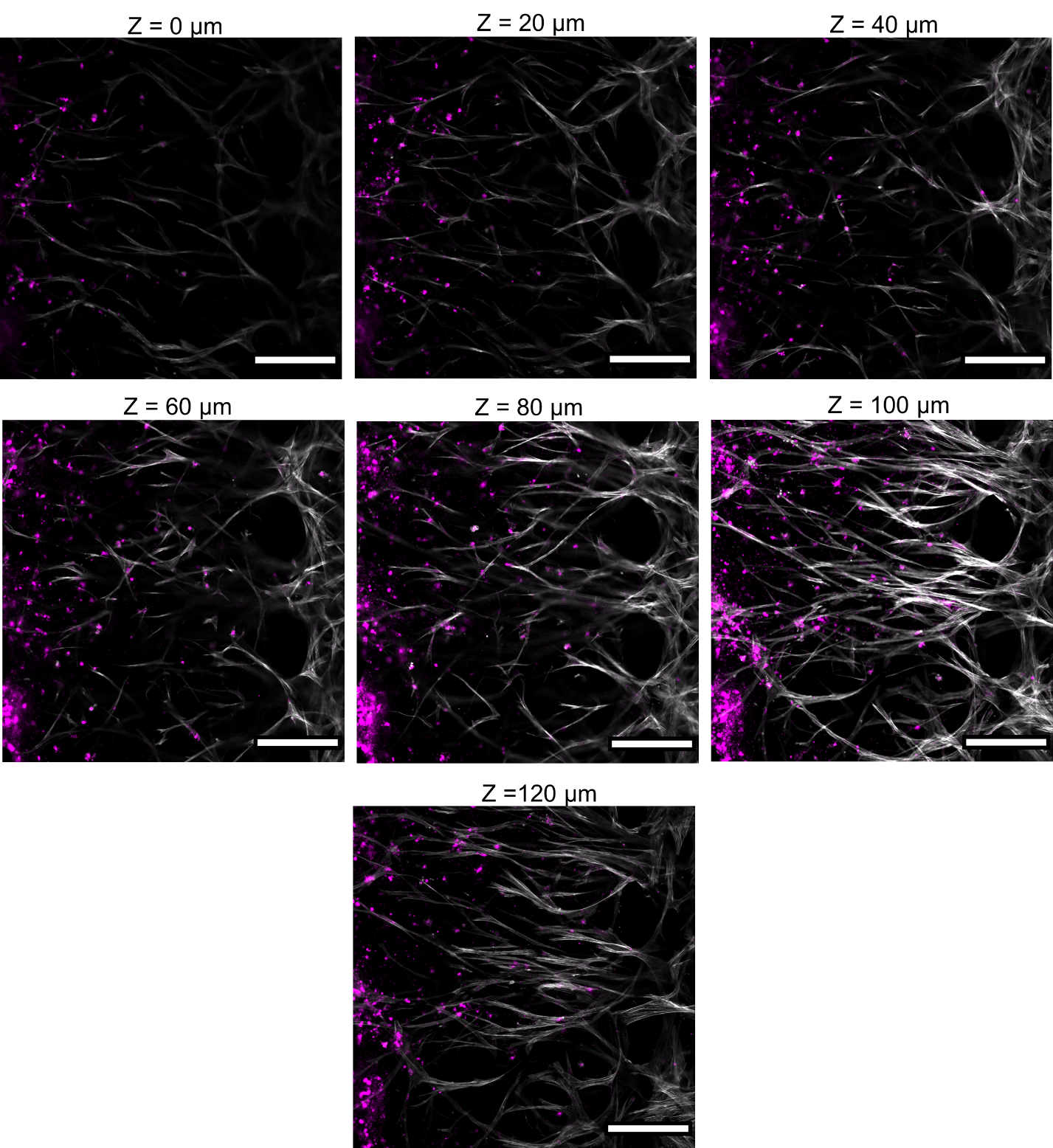

**Fig. S6. TNF-α-matured monocyte-derived dendritic cells (TNF-moDC)** **migration across Channel 4 in a FRC hydrogel.** Representative confocal microscopy images showing TNF-moDC migration across Channel 4 in different Z-stack planes at day 3. Magenta = CM-Dil labeled TNF-moDCs, white = phalloidin F-actin. Scale bars = 200 μm.

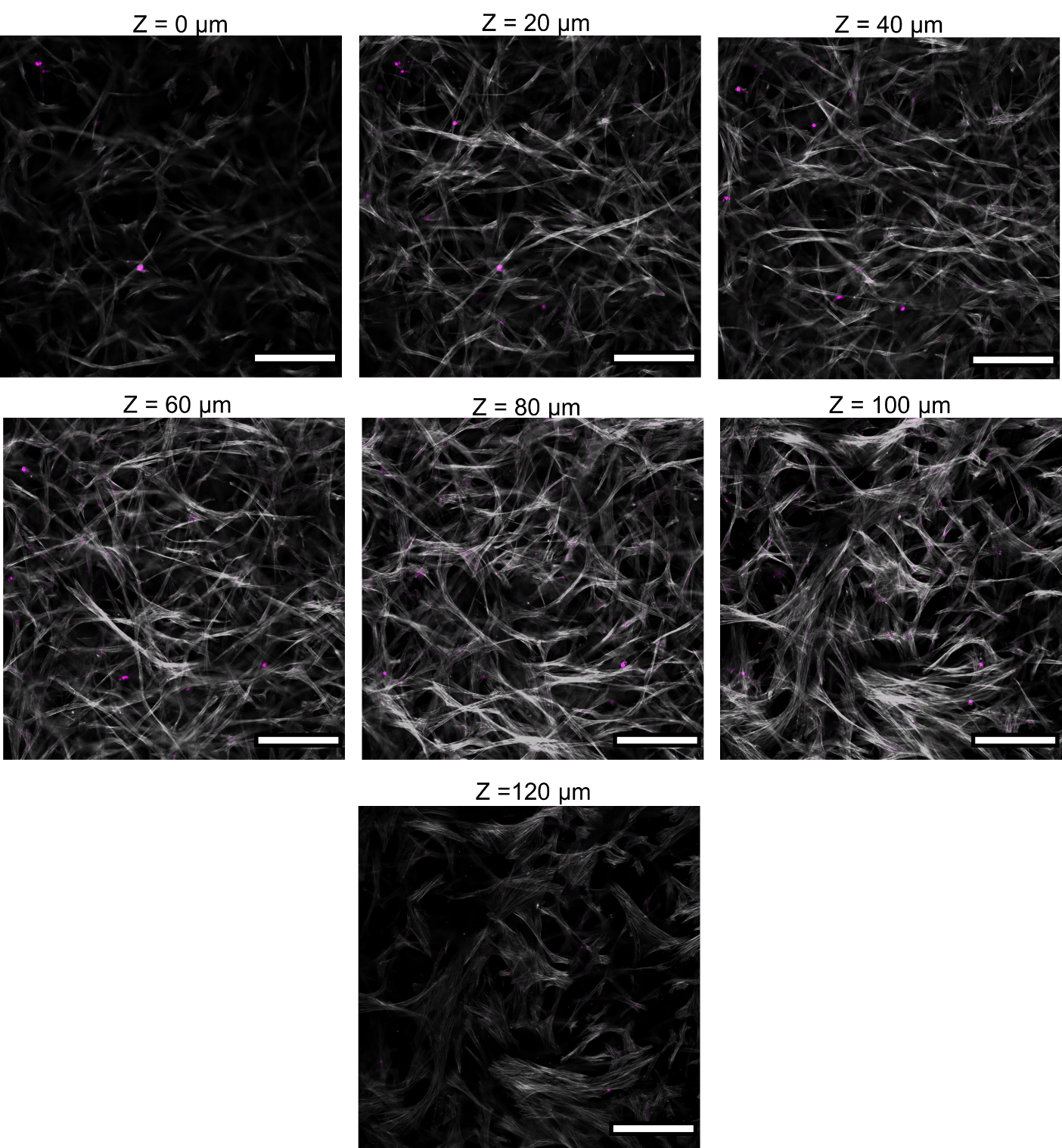

**Fig. S7. TNF-α-matured monocyte-derived dendritic cells (TNF-moDC) migration into Channel 5 in a FRC hydrogel.** Representative confocal microscopy images showing TNF-moDC migration into Channel 5 in different Z-stack planes at day 3. Magenta = CM-Dil labeled TNF-moDCs, white = phalloidin F-actin. Scale bars = 200 μm.

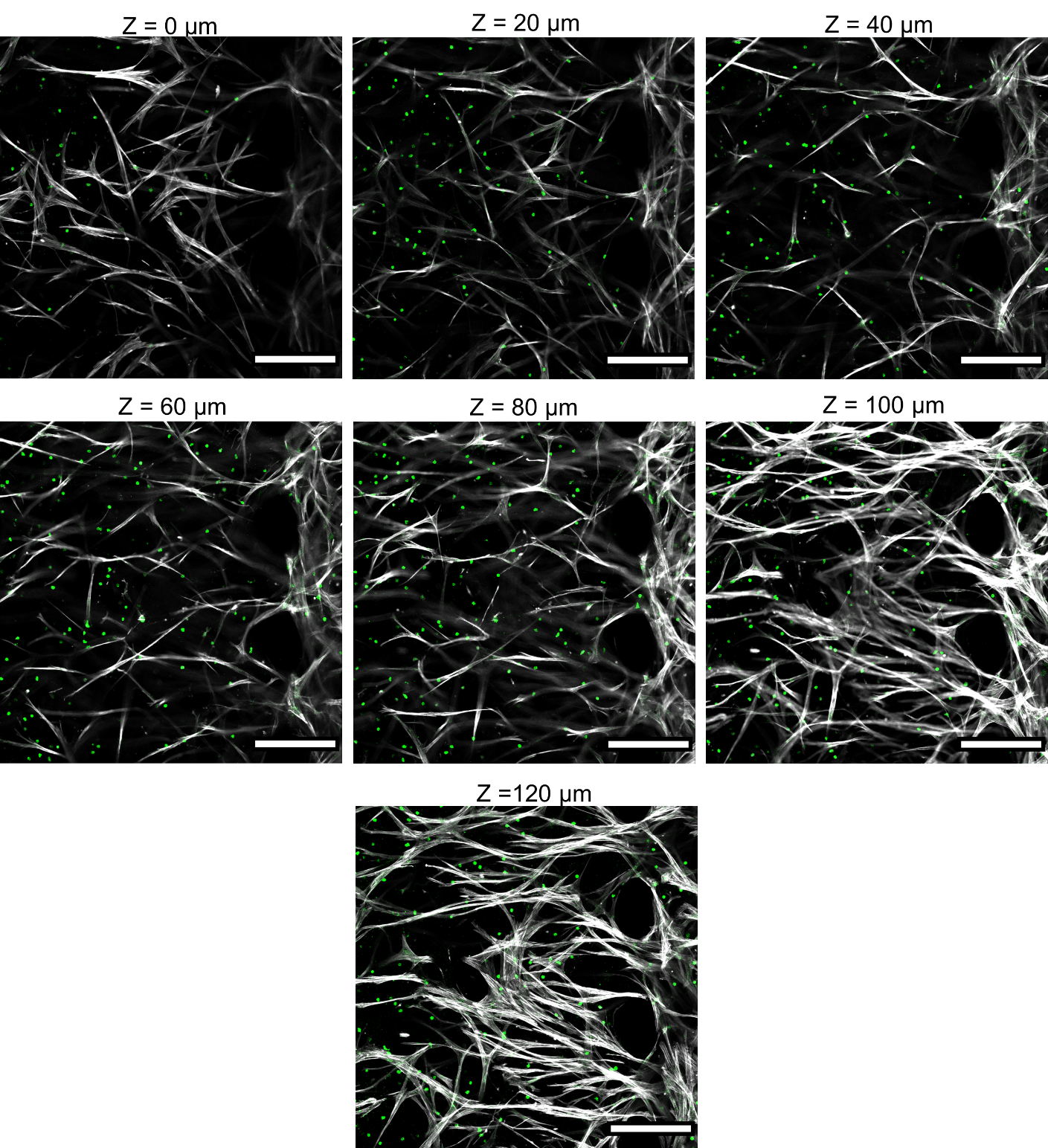

**Fig. S8. Pan T-cell migration across Channel 4 in a FRC hydrogel.** Representative confocal microscopy images showing pan T-cell migration across Channel 4 in different Z-stack planes at day 3. Green = CD3 labeled T-cells, white = phalloidin F-actin. Scale bar = 200 μm.

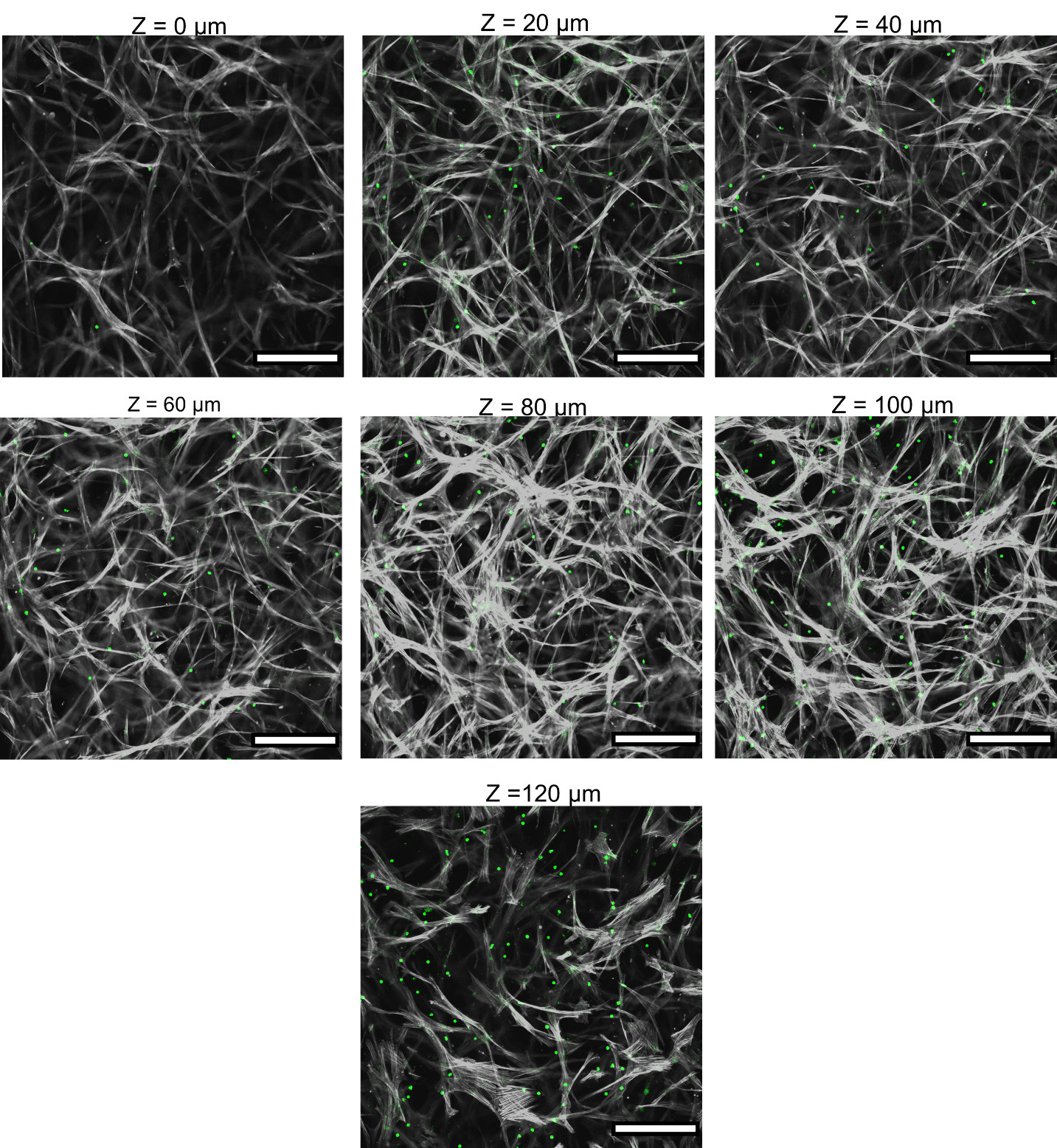

**Fig. S9. Pan T-cell migration into Channel 5 in a FRC hydrogel.** Representative confocal microscopy images showing pan T-cell migration into Channel 5 in different Z-stack planes at day 3. Green = CD3^+^ T-cells, white = phalloidin F-actin. Scale bar = 200 μm.

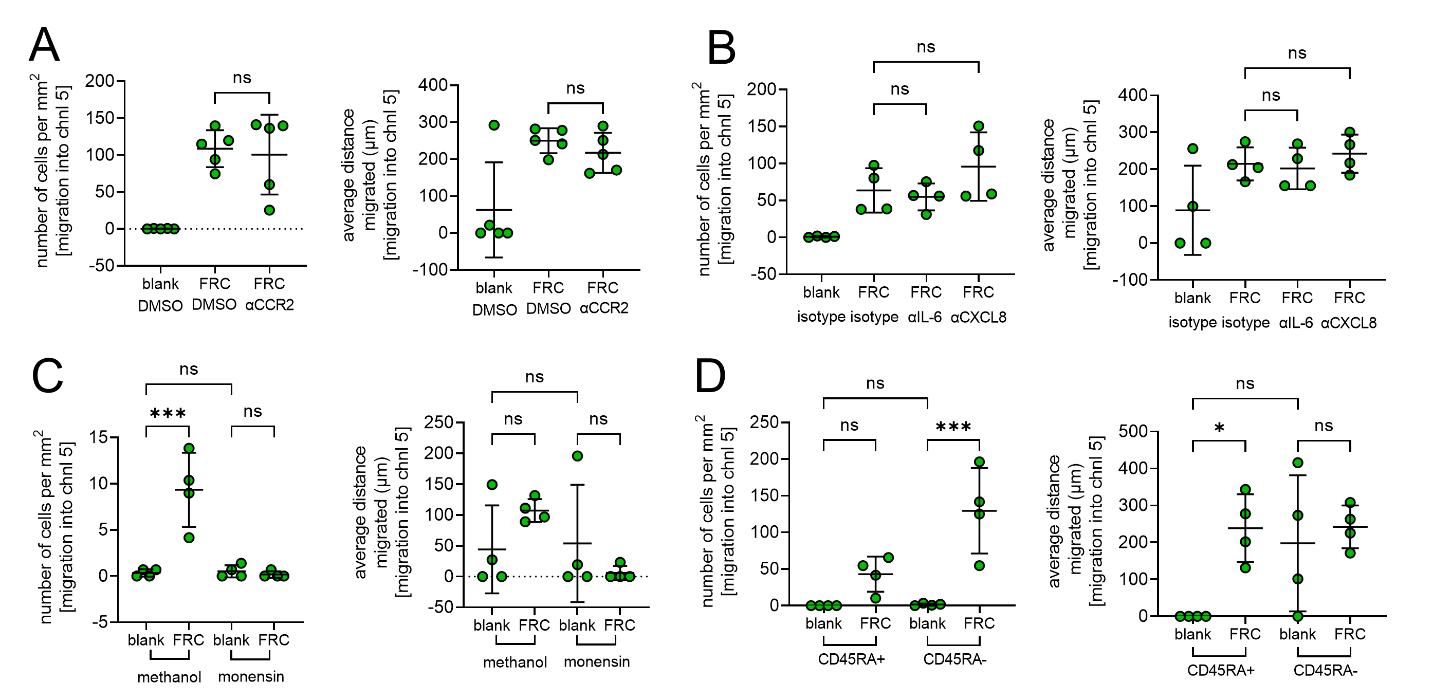

**Fig. S10. T-cell migration across lymph node stromal network chip.** (A) Quantification of pan T-cell migration into Channel 5 toward a blank hydrogel and a FRC hydrogel with or without a CCR2 antagonist. n = 5 devices. (B) Quantification of pan T-cell migration into Channel 5 toward a blank hydrogel and a FRC hydrogel in the presence of an isotype control antibody, anti-IL-6 antibody, and anti-CXCL8 antibody. n = 4 devices. (C) Quantification of pan T-cell migration into Channel 5 toward a blank hydrogel and a FRC hydrogel with or without monensin. n = 4 devices. (D) Quantification of CD45RA^+^ or CD45RA^-^ T-cell migration into Channel 5 toward a blank hydrogel and a FRC hydrogel. n = 4 devices. Data are presented as means ± SD. Significance is denoted by **P* < 0.05 or ****P* < 0.001 by one-way analysis of variance (ANOVA) with Tukey’s post hoc test.

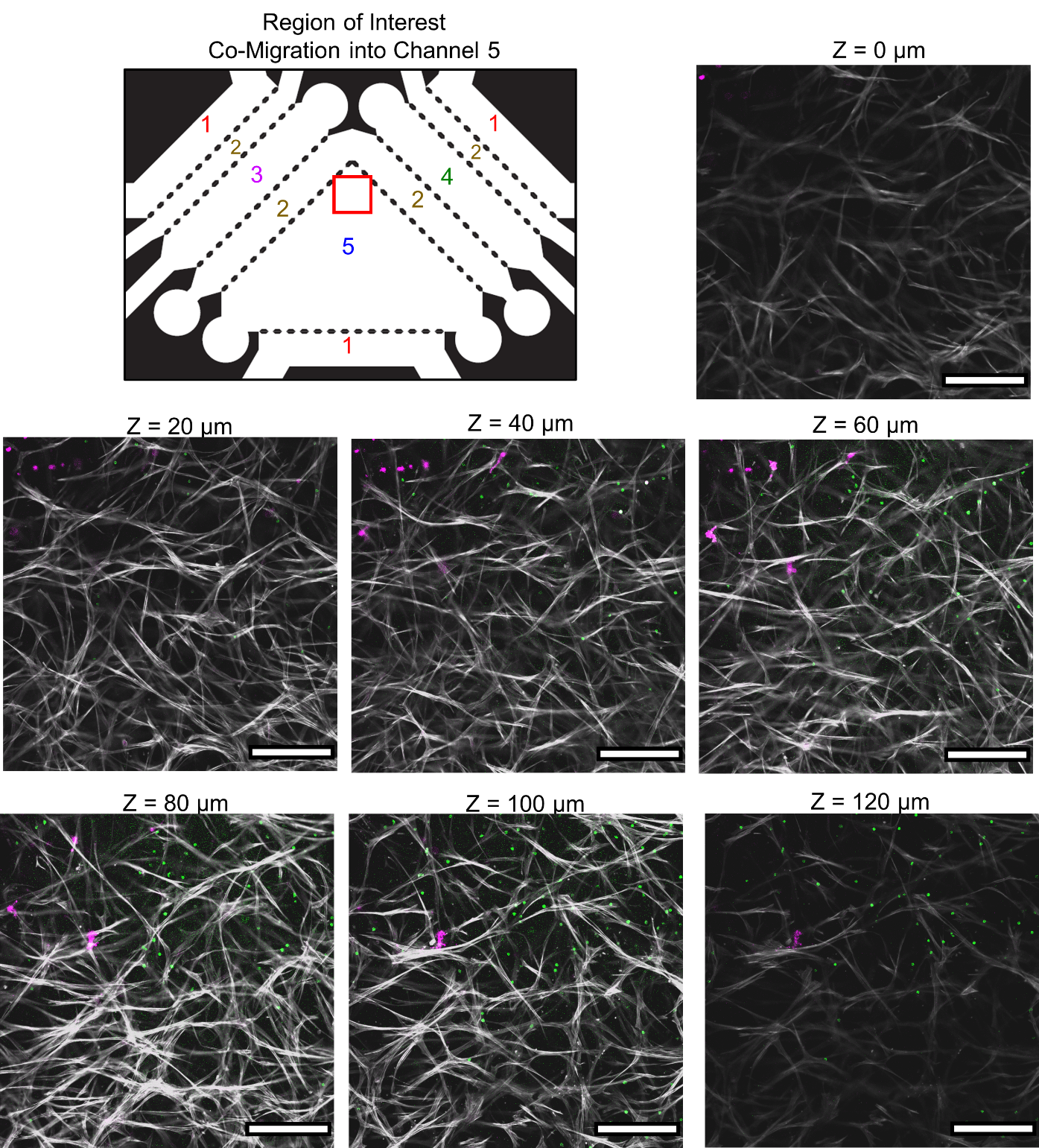

**Fig. S11.** **Migration of TNF-α-matured monocyte-derived dendritic cells (TNF-moDC) and pan T-cells into lymph node stromal network.** (Top left) Zoomed in image of photolithography mask denoting region in red box where confocal Z-stack images were taken across all devices and experiments to quantify migration into Channel 5. (Top right and bottom) Representative confocal microscopy images showing TNF-moDC and pan T-cell migration into Channel 5 in different Z-stack planes at day 3. Magenta = CM-Dil labeled dendritic cells, green = CD3 labeled T-cells, white = phalloidin F-actin. Scale bar = 200 μm.
